# Determination of suitable reference genes for RT-qPCR analysis in *Gryllodes sigillatus* (*Orthoptera*: *Gryllidae*)

**DOI:** 10.64898/2026.04.04.716481

**Authors:** Ben-Miled Houda, Romaric Bonhomme, Fanny Renois, Marie-Hélène Deschamps, Marie-Odile Benoit-Biancamano, François Meurens

## Abstract

The tropical house cricket *Gryllodes sigillatus* is a major species used in the edible insect farming industry. Despite the rapid expansion of this sector, diagnostic tools for detecting infections in these species remain limited. The lack of validated reference genes compromises the reliability of RT-qPCR-based gene expression analyses, which are essential for the development of molecular tools for disease diagnosis and health monitoring in insect production systems.

To address this gap, we evaluated the expression stability of six candidate reference genes (*ACTB, EF1, GAPDH, HisH3, RPL5*, and *18SrRNA*) across four body parts (abdomen, head, legs, and whole body) using a combination of complementary statistical approaches, including geNorm, NormFinder, BestKeeper, the ΔCt method, the R statistical environment, and the integrated RefFinder tool. Candidate genes were identified and annotated using the recently published *G. sigillatus* genome, through sequence comparisons with closely related insect species using BLAST and reciprocal BLAST analyses, multiple sequence alignments. All procedures complied with MIQE 2.0 guidelines to ensure methodological rigor and transparency.

The results showed that *ACTB, EF1, RPL5*, and *18SrRNA* exhibited stable and consistent expression across all analyzed tissues, whereas *GAPDH* and *HisH3* displayed high variability and were generally unsuitable for normalization, except in head tissue where *GAPDH* remained stable. This study provides the first validated set of reference genes for *G. sigillatus*, establishing a robust foundation for accurate, reproducible, and comparable gene expression analyses. Furthermore, these findings support the development of RT-qPCR-based diagnostic tools, contributing to improved health monitoring and biosafety in insect production systems.

## 1. Introduction

Insects have long been part of human consumption patterns and continue to hold substantial economic and cultural importance across multiple societies (Raheem et al., 2019). In the face of rapid global population growth and the concomitant increase in protein demand, intensive commercial insect farming is attracting growing interest as a sustainable alternative to conventional livestock systems (Huis, 2020). This emerging sector is fully aligned with the United Nations Sustainable Development Goals, particularly with regard to food security, reduction of environmental footprint, and efficient use of natural resources (J. D. Kong et al., 2025; UN, 2015).

In this context, research and development play a decisive role in ensuring efficient, safe, and sustainable growth of the edible insect industry. The production of robust scientific data is essential to guide, improve, and optimize farming practices, health management, and the selection of the most efficient species (J. D. Kong et al., 2025; Van Peer et al., 2024)

Among edible insects, crickets occupy a privileged position. They have been consumed by humans for thousands of years, and more than 60 species are currently integrated into food systems across a wide range of countries worldwide (Magara et al., 2021). In North America, two species dominate commercial production: the house cricket *Acheta domesticus* and the tropical cricket *Gryllodes sigillatus* (Larouche et al., 2023).

*Gryllodes sigillatus* currently appears as a particularly promising alternative to other commonly farmed species. This species exhibits high adaptive plasticity under varied ecological conditions, a relatively herbivorous diet, limited aggressiveness, as well as a more neutral odor characteristics that are advantageous for intensive farming systems (Magara et al., 2025; Tanga et al., 2025). Moreover, *G. sigillatus* shows excellent feed conversion efficiency, supporting its relevance for mass production systems for both human and animal food purposes (Alfiko et al., 2022; Hassan et al., 2024). Its rapid growth, strong reproductive potential, and good organoleptic qualities explain the growing interest it attracts in modern commercial farming systems (W. Kong et al., 2024; Magara et al., 2021).

Beyond their agronomic and industrial interest, crickets constitute model organisms widely used in fundamental research, particularly in evolutionary ecology, insect physiology, and immunology (Levy et al., 2024). They possess a functional immune system capable of responding effectively to various microbial stimuli through the activation of effector mechanisms such as lysozyme-type antimicrobial activity, melanization, encapsulation, and hemocyte mobilization (Duffield et al., 2022; Park & Stanley, 2015). However, while the physiological and cellular responses associated with infections have been described, the molecular bases of immunity in crickets remain insufficiently characterized. Reliable quantification of the expression of genes involved in these responses is essential for an integrated mechanistic understanding of immunity and for the development of molecular tools dedicated to health monitoring in farming systems (Duffield et al., 2022; Kingsolver et al., 2013).

Despite the growing economic importance of *G. sigillatus*, the genetic resources available for this species remain limited (Zhang et al., 2025), and no set of reference genes has yet undergone rigorous validation for RT-qPCR-based gene expression analyses. This methodological gap compromises the reliability of studies, particularly those focusing on immunity, for which gene expression may vary significantly among tissues, and the use of unvalidated constitutive genes is likely to introduce interpretative biases and lead to erroneous estimation of immune responses. In this context, the identification and validation of stable reference genes constitute an essential preliminary step for reliable analyses of immunity-related gene expression in *G. sigillatus*. This approach enables the establishment of a robust methodological framework for molecular studies of immune mechanisms and supporting the development of tools for both fundamental and applied research. In the longer term, such advances are indispensable for improving health management in farming systems, optimizing production practices, and supporting the sustainable and safe development of this emerging sector.

The objective of the present study was to identify and validate robust reference genes for RT-qPCR data normalization in *Gryllodes sigillatus*. To determine the most stable candidates, six genes commonly used in expression analysis beta-actin (*ACTB*), elongation factor 1 alpha (*EF1*), glyceraldehyde-3-phosphate dehydrogenase (*GAPDH*), histone H3 (*His H3*), ribosomal protein L5 (*RPL5*), and 18S ribosomal RNA (*18S rRNA*) were evaluated across four adult body parts (head, legs, abdomen, and whole body). Expression stability was analyzed using six complementary statistical approaches (geNorm, NormFinder, BestKeeper, ΔCt methods, RefFinder, and analyses in R), allowing cross-validation and rigorous assessment of each gene’s performance. Final validation of the selected genes was carried out by normalizing the expression of elongation factor 2 (*EF2*). This study thus establishes a solid methodological framework for gene expression analyses in *G. sigillatus* and supports the development of reliable molecular tools. It can be applied to health status assessment, early detection of immune imbalances, and optimization of cricket farming practices. It also contributes to infection monitoring, for which reference genes are essential for result normalization.

## 2. Materials and methods

### 2.1. Insects Samples

Adult tropical house crickets (*Gryllodes sigillatus*) individuals were obtained from a local pet store and selected based on their healthy condition. No individuals exhibited visible signs of physiological alteration, such as wing abnormalities, external lesions, or abnormal mortality. Specimens were transported to the laboratory in appropriate ventilated containers and immediately stored at −80 °C until RNA extraction procedures.

### 2.2. Total RNA Extraction and Reverse Transcription

Adult tropical house crickets (*G. sigillatus*) were used. Abdomen, head, and leg tissues, as well as whole-body samples, were dissected separately, with antennae removed from head samples. Each sample type consisted of ten independent biological replicates.

Total RNA was extracted individually using the RNeasy Plus Mini Kit (Qiagen, Stockach, Germany) according to the manufacturer’s instructions, including an on-column DNase treatment to eliminate genomic DNA contamination. Tissues were homogenized in RLT buffer containing proteinase K and incubated at 60 °C for 6 h. RNA was eluted in 30 µL of RNase-free water.

RNA quantity and purity were assessed by spectrophotometry using a NanoDrop™ One/OneC (Thermo Fisher Scientific). cDNA synthesis was performed using 500 ng of total RNA per sample with the iScript™ Advanced cDNA Synthesis Kit for RT-qPCR (Bio-Rad, Hercules, CA, USA), following the manufacturer’s recommendations. Reverse transcription was carried out under the following conditions: 5 min at 25 °C, 20 min at 46 °C, and 1 min at 95 °C. No–reverse transcriptase (–RT) controls were included for each sample type.

The resulting cDNA was then adjusted to a final volume of 40 µL for RT-qPCR analyses. Total RNA was stored at −80 °C, whereas cDNA was stored at −20 °C until use.

### 2.3. Identification of Novel Candidate Reference Genes

Candidate gene selection relied on the *Gryllodes sigillatus* reference genome available in NCBI. (https://www.ncbi.nlm.nih.gov/), as well as on the recently published complete reference genome (GenBank: GCA_965111905.1) (Zhang et al., 2025). This approach was complemented by the inclusion of reference genes commonly used in other insect species, as reported in previous studies (Lü et al., 2018).

Because the *G. sigillatus* reference genome has been only recently published and remains poorly annotated, sequence identification focused on retrieving homologs of the candidate reference genes described above from better-characterized insect species. To this end, BLASTx and reciprocal tBLASTn analyses were performed to identify expressed sequencescorresponding to these reference gene homologs in *G. sigillatus*. BLASTx searches were conducted using nucleotide sequences of candidate reference genes from phylogenetically related insects, particularly other cricket species such as *Acheta domesticus* and *Gryllus bimaculatus*, as well as *Locusta migratoria* (Foquet & Song, 2020; Y. Wang et al., 2007; Q. Yang et al., 2014), in order to identify conserved homologous protein regions. Reciprocal tBLASTn analyses were carried out using reference protein sequences corresponding to these genes from model insects, particularly *Sphingonotus tsinlingensis* (L. Zhao et al., 2021), to query the available transcript and EST databases for *G. sigillatus*, following approaches previously described in other animal species (Bruel et al., 2010).

The sequences corresponding to genes of interest were subsequently selected for the design of gene-specific primers for RT-qPCR analyses, in accordance with MIQE 2.0 guidelines (Bustin et al., 2009, 2025).

### 2.4. Primer Design and evaluation

Twelve primer pairs were designed from the coding sequences (CDS) of the target genes using Clone Manager 9 software (Scientific & Educational Software, Cary, NC, USA), and one additional primer pair was selected from the literature (Table 1). Primer design followed MIQE guidelines (Bustin et al., 2009, 2025), including primer lengths of 18–20 nucleotides, annealing temperatures of 55–59 °C, and amplicon sizes ranging from 225 to 326 bp. Primers were synthesized by Integrated DNA Technologies (IDT, Ontario, Canada).

**Table 1:**
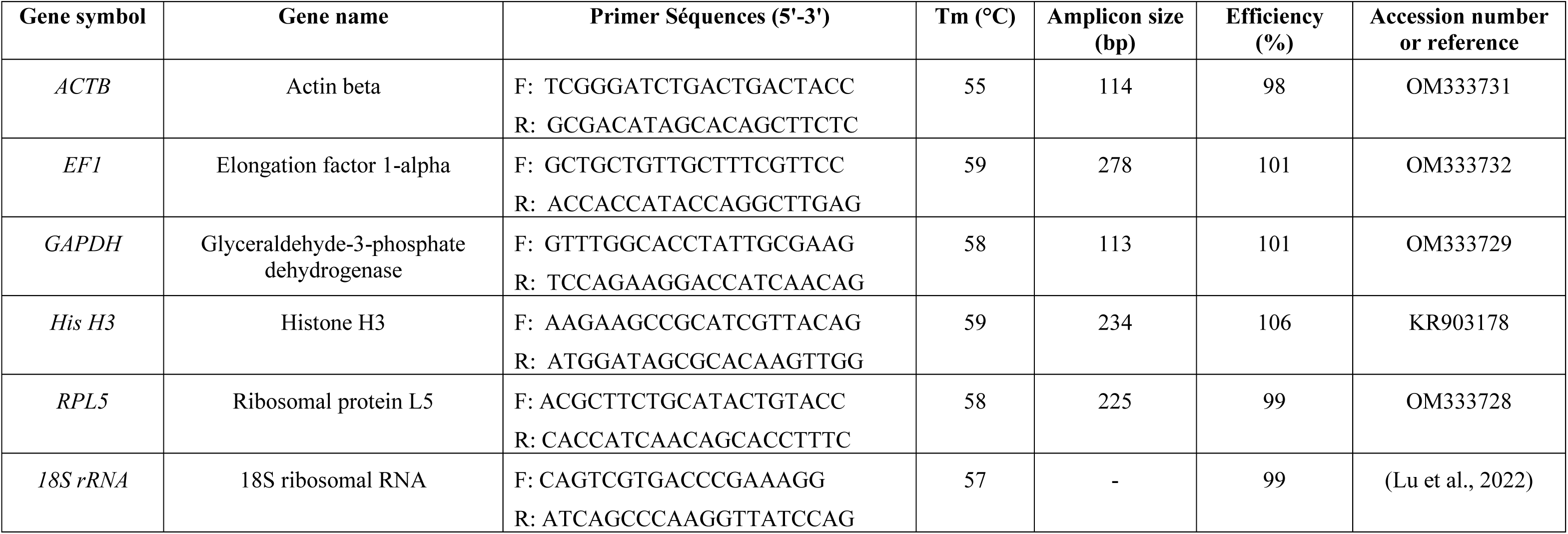
Primer specificity and amplification characteristics of candidate reference genes.

Primer specificity was validated by qPCR using cDNA as the template. Amplification efficiencies were determined from standard curves generated using serial dilutions of a pooled cDNA sample. Depending on the analysis, standardized 1:2 serial dilutions were performed, and quantification cycle (Cq)values were plotted against the corresponding dilution factors. The slope of each standard curve was determined by linear regression, and PCR efficiency was calculated using the equation E (%) = (10^^(–1/slope)^ – 1) × 100. Six primer pairs were selected for subsequent analyses, corresponding to the reference genes *ACTB*, *EF1*, *GAPDH*, *His H3*, *RPL5*, and *18S rRNA*. Primer sequences, amplicon sizes, and amplification efficiencies are reported in Table 1.

### 2.5. Reverse Transcription Quantitative Polymerase Chain Reaction (RT-qPCR)

All RT-qPCR reactions were performed using a CFX Opus 96 real-time PCR system (Bio-Rad, Hercules, USA) with SsoAdvanced Universal SYBR Green Supermix chemistry. Each reaction was prepared in a final volume of 10 µL containing 2 µL of previously synthesized cDNA diluted 1:10, 5 µL of 2× Supermix (Bio-Rad, Hercules, CA, USA), 0.2 µL of each primer (forward and reverse; IDT, Ontario, Canada), and nuclease-free water to adjust the final volume. Analyses were conducted using ten biological replicates with two technical replicates per sample. No-template controls (NTCs) were systematically included to monitor potential contamination.

The thermal cycling program consisted of an initial activation at 98 °C for 30 s, followed by 40 cycles at 98 °C for 5 s and 55–59 °C for 30 s. Amplification specificity was verified by melt curve analysis (65–95 °C, 0.5 °C increments). Quantification cycle (Cq) values were determined using CFX Maestro software (v2.3) with a single threshold. All real-time qPCR analyses were performed in accordance with the MIQE guidelines (Bustin et al., 2009, 2025).

### 2.6. Stability analysis

The expression stability of six candidate reference genes was evaluated using RefFinder (Xie et al., 2012; https://www.ciidirsinaloa.com.mx/RefFinder-master/), which integrates the geNorm (v3.5; Vandesompele et al., 2002), NormFinder (v0.953) (Andersen et al., 2004), BestKeeper (v1) (Pfaffl et al., 2004), and DeltaCt (Cycle threshold) method (Silver et al., 2006). The geNorm calculates the expression stability value (M) (Vandesompele et al., 2002), with genes exhibiting lower M values considered more stable; an M value < 1 is generally accepted as the threshold for classifying genes as relatively stable (Hellemans et al., 2007; Kim & Kim, 2025; Liu et al., 2014).

NormFinder estimates both intra- and inter-group expression stability, while BestKeeper identifies stable genes based on low variation in Cq values (standard deviation, SD < 1) (Chechi et al., 2012; Jeon et al., 2020). The Delta Ct method compares the mean Cq differences between pairs of genes. Pairwise variation analysis (Vₙ/Vₙ₊₁) was performed using R software (v4.5) (*R Core Team 2021*) to determine the optimal number of reference genes required for normalization, applying a cut-off value of <0.15, as recommended by Vandesompele and collaborators (2002) (Hellemans & Vandesompele, 2014; Vandesompele et al., 2002).

### 2.7. Validation of reference gene selection

The *EF2* (elongation factor 2) gene was used as a target gene to assess the impact of reference gene selection on the normalization of RT-qPCR data. *EF2* expression levels were measured in different tissues and normalized using the selected candidate reference genes (Kaul et al., 2011). EF2-specific primers were designed using Clone Manager 9 software (Scientific & Educational Software, Cary, NC, USA). The primer sequences were as follows: F: GGCTGGAGAAGCTTTACTAC and R: CATCATGAGGCCCTTCATAC, generating an amplicon of 101 bp. The annealing temperature was set at 56 °C.

Relative expression of *EF2* was calculated using the 2-ΔΔCq method (Livak & Schmittgen, 2001). For each tissue, normalization was performed using pairs of reference genes identified as the most stable, as well as using pairs of unstable reference genes, in order to evaluate the impact of reference gene selection on the robustness of normalization. Comparisons between the two normalization strategies (stable vs unstable reference genes) were performed using paired statistical tests, to account for the fact that the same samples were analyzed using both normalization methods. Data normality was assessed using the Shapiro–Wilk test and was confirmed for all analyses. Consequently, comparisons were conducted using paired two-tailed Student’s t-tests. Statistical analyses were performed using GraphPad Prism 8 software (GraphPad, San Diego, CA, USA), and a p value < 0.05 was considered statistically significant.

## 3. Results

### 3.1. Primer Specificity Validation

The optimal annealing temperature and specificity of the primer pairs were evaluated using pooled cDNA, prepared separately for each tissue type from the same reverse transcription reaction, with equal template quantities per sample. A temperature gradient ranging from 55 to 60 °C was applied to identify the annealing temperature that ensured specific and efficient amplification for each primer pair (Table 1).

Amplification specificity was confirmed by melting curve analysis, which revealed a single peak for each gene (Figure 1), with no evidence of primer-dimer formation. No amplification was detected in the no-template negative controls (NTC).

**Figure 1:**
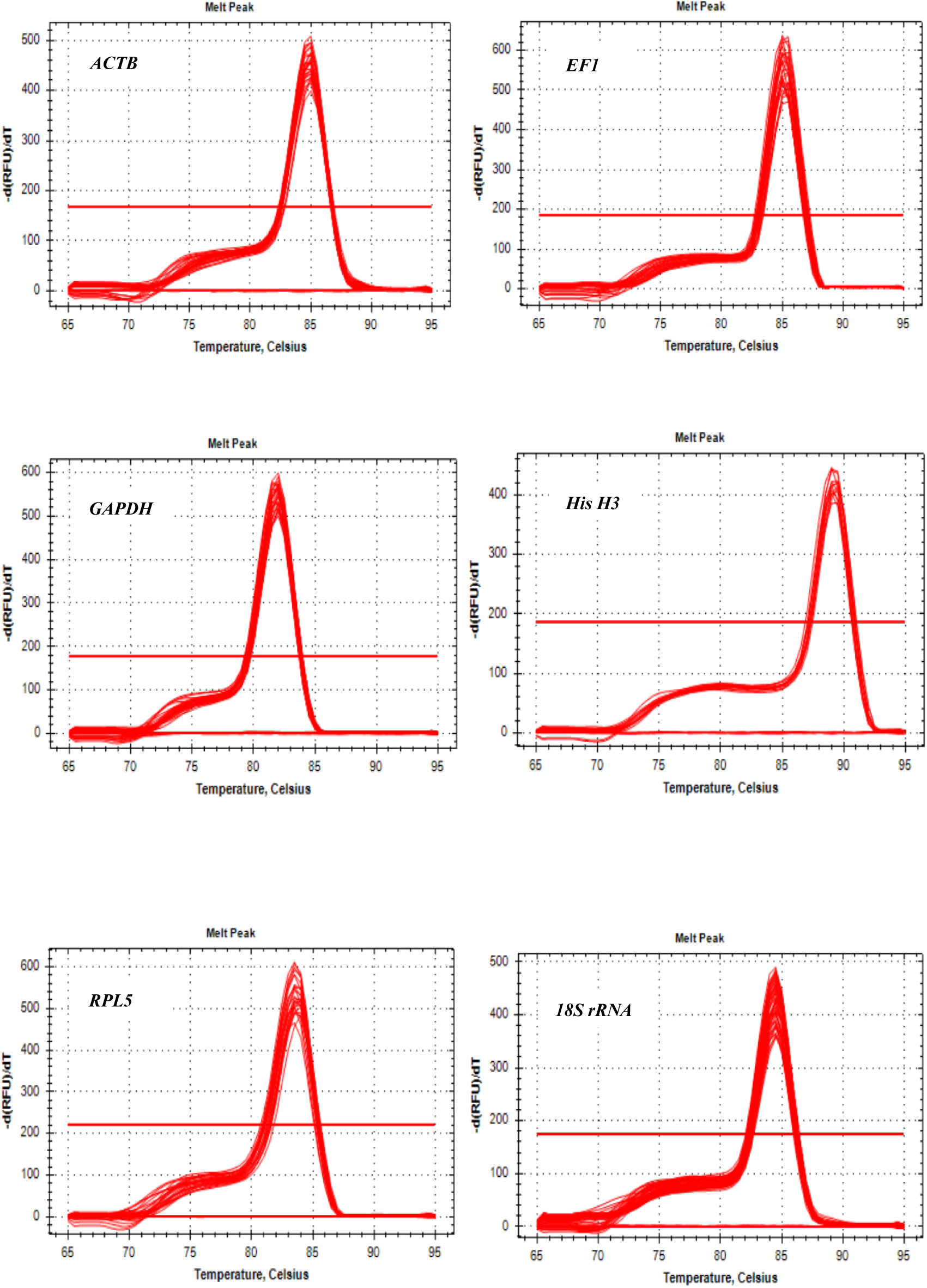
Melting curve profiles of the six candidate reference genes in *Gryllodes sigillatus*

The amplification efficiencies (E) of the primer pairs ranged from 98% for *ACTB* to 106% for *His H3*, falling within the recommended range of 90–110%, while the correlation coefficients (R²) of the standard curves were all greater than 0.991. Based on these results, all primer pairs were retained for subsequent RT-qPCR analyses. Detailed information on the candidate reference genes, including primer sequences, amplicon lengths, and annealing temperatures, is provided in Table 1.

### 3.2. Expression levels of candidate reference genes

The violin plot (Figure 2) integrates a central summary of the data, where the median is represented by a black dot, with a smooth density profile that depicts how values are distributed across the range. The lateral expansion of the shape corresponds to the relative concentration of observations at each level. In contrast to a box plot, this representation captures subtle features of the data distribution and offers a more comprehensive view of variability (Hintze & Nelson, 1998).

**Figure 2:**
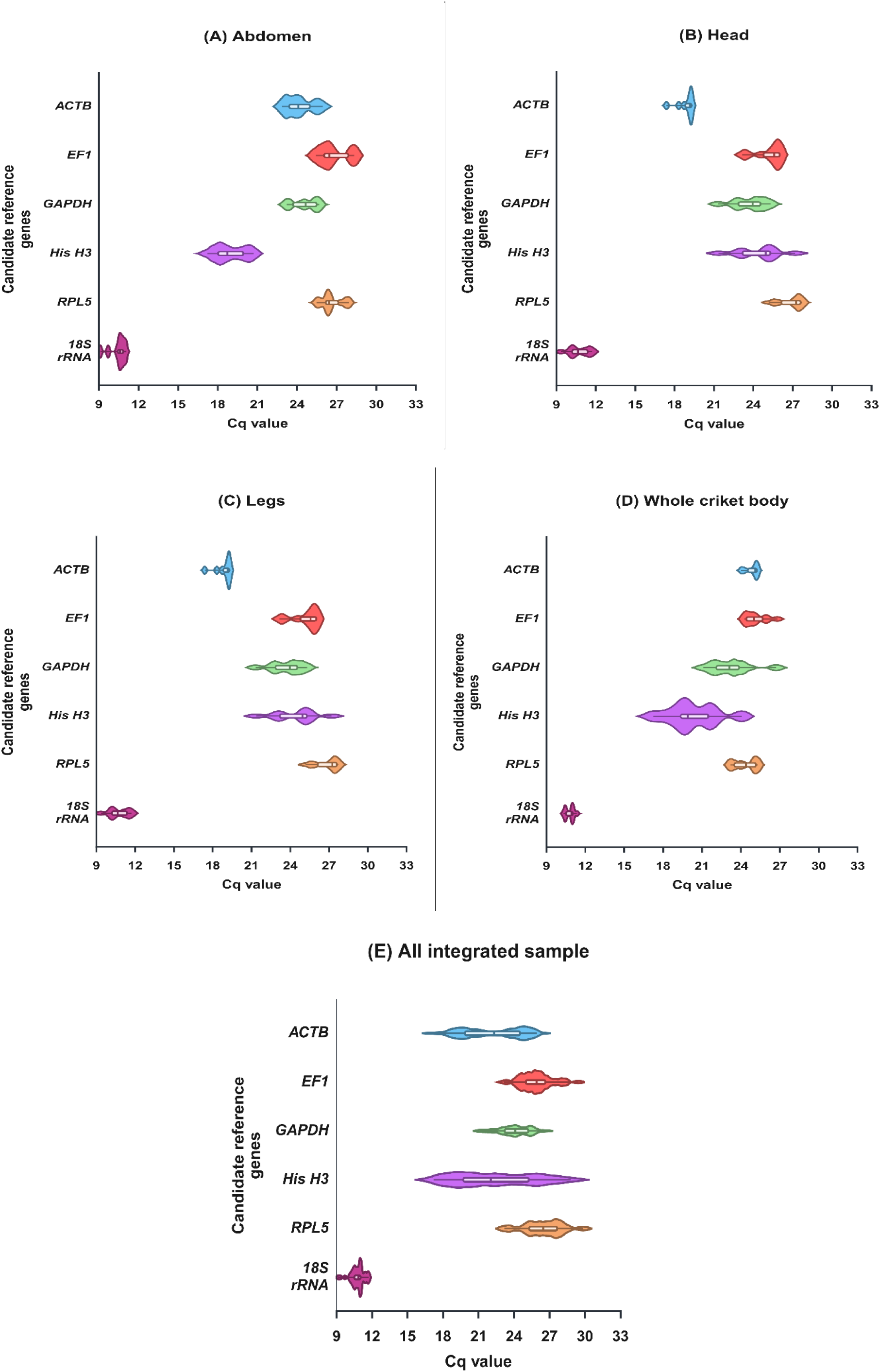
Violin plot representing the distribution of Cq values for *ACTB, EF1, GAPDH, HisH3, RPL5*, and *18S rRNA* across four tissues (abdomen, head, legs, and whole body) of the tropical cricket *Gryllodes sigillatus* and all integrated samples. The colored dot in each violin plot represents the median Cq value, while the white bar indicates the interquartile range. The width of the violin reflects the data density at a given horizontal coordinate (frequency of values along the x-axis). The different colors in the violin plots represent the various candidate reference genes.

The raw Cq values directly reflect gene expression levels and constitute essential baseline data for the evaluation of candidate reference genes (Chen et al., 2015; X. Tang et al., 2017). Gene expression is inversely proportional to the Cq value, with lower Cq values indicating higher expression levels, and vice versa. In this study, RT-qPCR was used to analyze the expression profiles of six candidate reference genes across different tissues.

As shown in Figure 2, the mean Cq values of the six reference genes ranged from 10.47 (*18S rRNA*) to 26.89 (*RPL5*) among the different tissues analyzed, and from 10.77 (*18S rRNA*) to 26.47 (*RPL5*) when considering all samples. Most Cq values were distributed between 19 and 27. Analysis of Cq value distributions indicated that gene expression varied depending on the tissue type.

Across all tissues examined, *18S rRNA* consistently exhibited the lowest Cq values, clustered around 10, indicating that it was the most abundantly expressed gene among the candidates evaluated. In contrast, the other reference genes showed comparable expression levels across tissues, with low variation in Cq values, suggesting relatively homogeneous expression.

### 3.3 Expression stability analysis of candidate reference genes using geNorm

Expression stability of candidate reference genes was evaluated using the geNorm algorithm, applied independently to each tissue and to the combined dataset including all tissues. geNorm ranks candidate reference genes based on the average expression stability value (M), allowing genes to be ordered from the most stable to the least stable (Figure 3). In accordance with previous studies, a mean stability value of M < 1 was considered an acceptable threshold to classify genes as relatively stable (Hellemans & Vandesompele, 2014; Kim & Kim, 2025; Liu et al., 2014).

**Figure 3:**
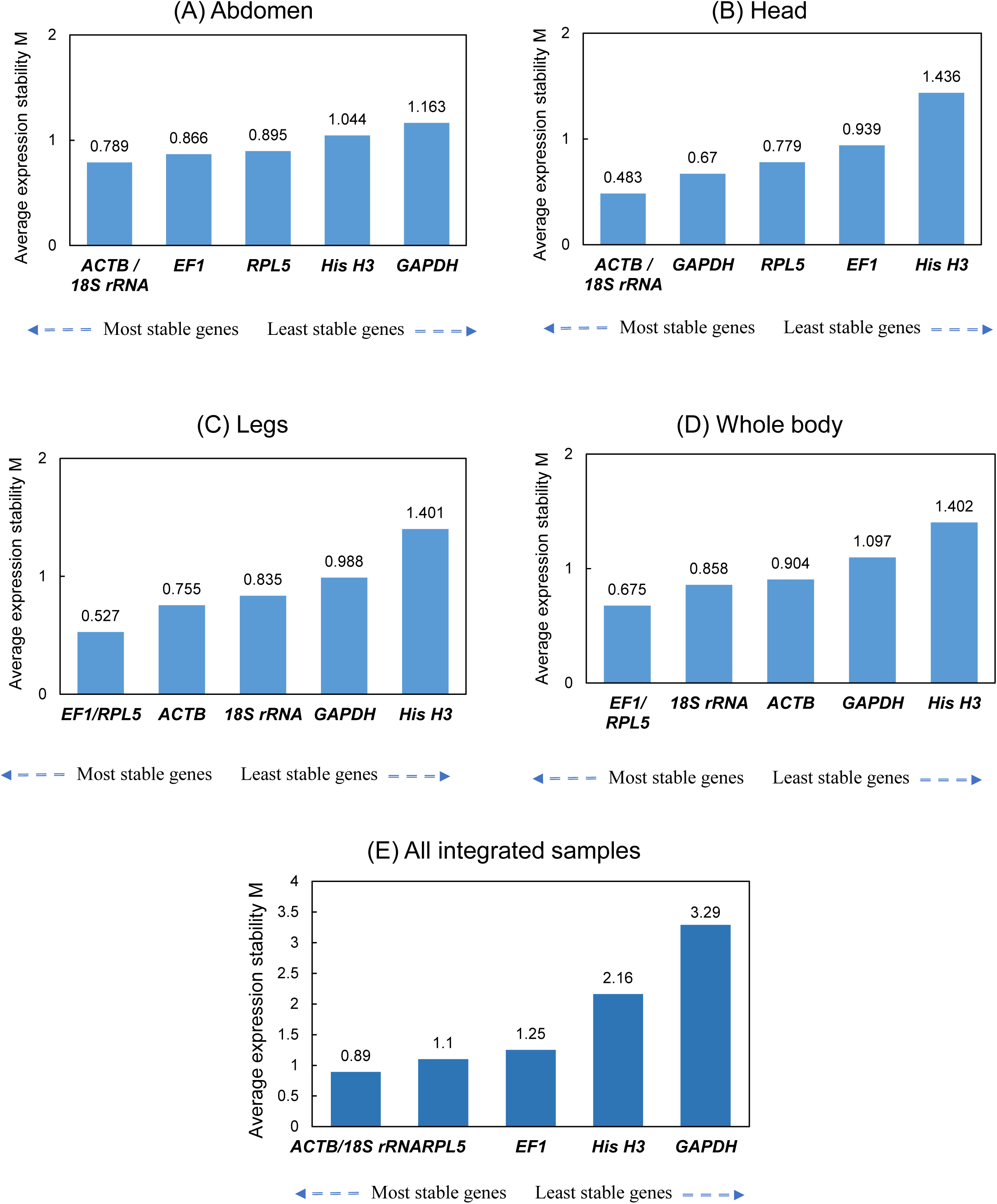
The expression stability values (M) of the six candidate reference genes in the different tissues of tropical cricket (*Gryllodes sigillatus*) and all integrated samples were assessed using the geNorm program. The least stable genes, displaying the highest M values, are located on the right side, whereas the most stable genes, with the lowest M values, are shown on the left.

In *G. sigillatus*, analysis of abdominal tissue showed that *ACTB*/*18S rRNA* exhibited the lowest stability values (M = 0.78), indicating relatively stable expression, whereas *GAPDH* was identified as the least stable gene, displaying the highest M value (M = 1.16) (Figure 3A). Comparable results were observed in head tissue, where *ACTB*/*18S rRNA* again demonstrated the highest expression stability (M = 0.48) (Figure 3 C). In contrast, *His H3* was identified as the least stable gene in this tissue, with the highest M value (M = 1.43).

For leg tissue and whole-body samples, *EF1*/*RPL5* showed the lowest stability values, with M values of 0.52 and 0.67, respectively. In both tissues, *GAPDH* consistently emerged as the least stable gene, reaching a high M value of 1.40 (Figure 3 B, D).

For all tissues analyzed individually, a threshold of M < 1 was applied. In contrast, for the analysis of the integrated tissues, characterized by heterogeneous values derived from different tissues, a more permissive threshold (M < 1.5) was adopted (Jeon et al., 2020). Applying this threshold to the combined dataset, *18S rRNA* and *ACTB* were identified as the most stable genes, with an M value of 0.89. They were followed by *RPL5* and *EF1*, with M values of 1.10 and 1.25, respectively. Conversely, *HisH3* and *GAPDH* were considered unstable, as their M values exceeded 1.5 (Figure 3 E).

Overall, the geNorm analysis indicates that *GAPDH* and *His H3* are among the least stable reference genes across most analyzed tissues as well as in the combined dataset. A notable exception was observed in head tissue, where *GAPDH* showed acceptable stability with an M value below 1 (M = 0.67). These findings highlight the importance of tissue-specific validation of reference genes prior to RT-qPCR data normalization in *G. sigillatus*.

### 3.4. Expression Stability Analysis by NormFinder

The NormFinder analysis, which accounts for both intra-group and inter-group variation, was used to evaluate the stability of candidate reference genes. Genes exhibiting the lowest stability values (SV) were considered the most stable (Pfaffl et al., 2004); however, no specific threshold value is recommended by this algorithm. The Figure 4 and Table 2 present the stability values of reference genes for each tissue type.

**Figure 4:**
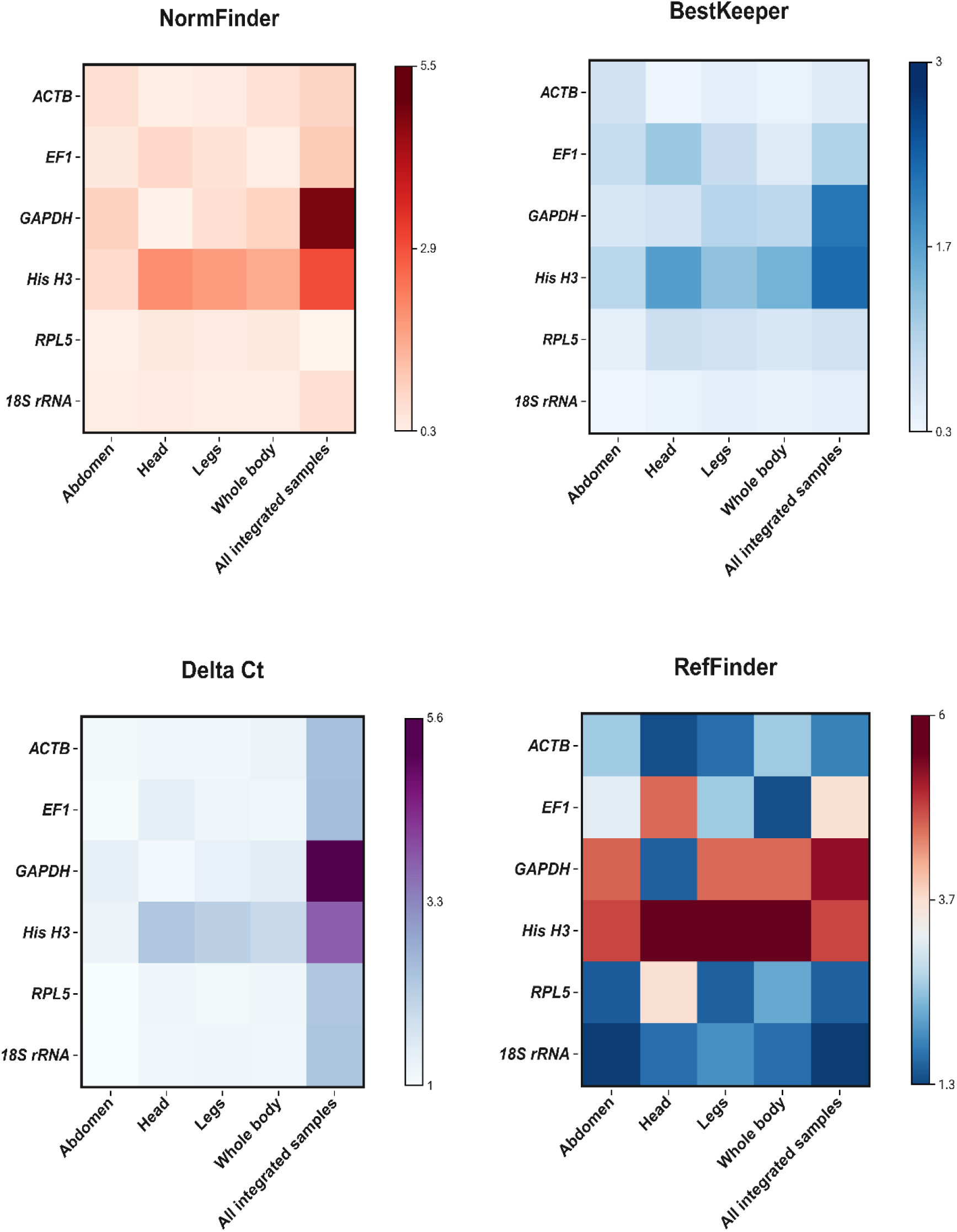
Stability analysis of the six candidate reference genes by NormFinder, BestKeeper, comparative ΔCt and RefFinder. Within a panel, the smaller the value, the lighter the colour, and the higher the stability of the candidate reference genes

**Table 2:** Stability ranking of the six candidate reference genes used in the present study according to different algorithms applied to four distinct tissues of the tropical cricket (*Gryllodes sigillatus*) and all integrated samples.

|  |  | NormFinder |  | BestKeeper |  | geNorm |  | Delta Ct |  | RefFinder |  |
| --- | --- | --- | --- | --- | --- | --- | --- | --- | --- | --- | --- |
|  | Rank | Gene | Stability | Gene | Stability | Gene | Stability | Gene | Stability | Gene | Stability |
| **Abdomen** | 1 | *RPL5* | 0.49 | *18S rRNA* | 0.40 | *18S rRNA* | 0.78 | *RPL5* | 1.01 | *18S rRNA* | 1.41 |
|  | 2 | *18S rRNA* | 0.57 | *RPL5* | 0.54 | *ACTB* | 0.78 | *18S rRNA* | 1.02 | *RPL5* | 1.68 |
|  | 3 | *EF1* | 0.68 | *GAPDH* | 0.72 | *EF1* | 0.86 | *EF1* | 1.08 | *ACTB* | 2.82 |
|  | 4 | *ACTB* | 0.88 | *ACTB* | 0.84 | *RPL5* | 0.89 | *ACTB* | 1.16 | *EF1* | 3.40 |
|  | 5 | *His H3* | 1.00 | *EF1* | 0.96 | *HisH3* | 1.04 | *His H3* | 1.28 | *GAPDH* | 5.04 |
|  | 6 | *GAPDH* | 1.18 | *His H3* | 1.08 | *GAPDH* | 1.16 | *GAPDH* | 1.40 | *His H3* | 5.23 |
| **Head** | 1 | *GAPDH* | 0.40 | *ACTB* | 0.42 | *ACTB* | 0.48 | *GAPDH* | 1.13 | *ACTB* | 1.56 |
|  | 2 | *ACTB* | 0.51 | *18S rRNA* | 0.48 | *18S rRNA* | 0.48 | *18S rRNA* | 1.18 | *GAPDH* | 1.73 |
|  | 3 | *18S rRNA* | 0.63 | *GAPDH* | 0.80 | *GAPDH* | 0.67 | *ACTB* | 1.19 | *18S rRNA* | 1.86 |
|  | 4 | *RPL5* | 0.67 | *RPL5* | 0.88 | *RPL5* | 0.77 | *RPL5* | 1.21 | *RPL5* | 4 |
|  | 5 | *EF1* | 1.06 | *EF1* | 1.32 | *EF1* | 0.93 | *EF1* | 1.45 | *EF1* | 5 |
|  | 6 | *HisH3* | 2.33 | *HisH3* | 1.84 | *His H3* | 1.43 | *His H3* | 2.43 | *His H3* | 6 |
| **Legs** | 1 | *18S rRNA* | 0.58 | *ACTB* | 0.56 | *EF1* | 0.52 | *RPL5* | 1.16 | *RPL5* | 1.73 |
|  | 2 | *ACTB* | 0.59 | *18S rRNA* | 0.56 | *RPL5* | 0.52 | *ACTB* | 1.17 | *ACTB* | 1.86 |
|  | 3 | *RPL5* | 0.64 | *RPL5* | 0.84 | *ACTB* | 0.75 | *18S rRNA* | 1.20 | *18S rRNA* | 2.21 |
|  | 4 | *EF1* | 0.83 | *EF1* | 0.94 | *18S rRNA* | 0.83 | *EF1* | 1.24 | *EF1* | 2.82 |
|  | 5 | *GAPDH* | 0.88 | *GAPDH* | 1.10 | *GAPDH* | 0.98 | *GAPDH* | 1.38 | *GAPDH* | 5 |
|  | 6 | *His H3* | 2.11 | *His H3* | 1.40 | *His H3* | 1.40 | *His H3* | 2.22 | *His H3* | 6 |
| **Whole cricket body** | 1 | *18S rRNA* | 0.53 | *ACTB* | 0.48 | *EF1* | 0.67 | *EF1* | 1.17 | *EF1* | 1.56 |
|  | 2 | *EF1* | 0.57 | *18S rRNA* | 0.50 | *RPL5* | 0.67 | *18S rRNA* | 1.18 | *18S rRNA* | 1.86 |
|  | 3 | *RPL5* | 0.67 | *EF1* | 0.64 | *18S rRNA* | 0.85 | *RPL5* | 1.21 | *RPL5* | 2.44 |
|  | 4 | *ACTB* | 0.87 | *RPL5* | 0.72 | *ACTB* | 0.90 | *ACTB* | 1.30 | *ACTB* | 2.82 |
|  | 5 | *GAPDH* | 1.17 | *GAPDH* | 1.04 | *GAPDH* | 1.09 | *GAPDH* | 1.52 | *GAPDH* | 5 |
|  | 6 | *His H3* | 1.85 | *His H3* | 1.60 | *His H3* | 1.40 | *His H3* | 2.01 | *His H3* | 6 |
| **All integrated samples** | 1 | *RPL5* | 0.34 | *18S rRNA* | 0.57 | *18S rRNA* | 0.89 | *RPL5* | 2.43 | *18S rRNA* | 1.41 |
|  | 2 | *18S rRNA* | 0.88 | *ACTB* | 0.63 | *ACTB* | 0.89 | *18S rRNA* | 2.49 | *RPL5* | 1.73 |
|  | 3 | *ACTB* | 1.12 | *RPL5* | 0.83 | *RPL5* | 1.10 | *ACTB* | 2.59 | *ACTB* | 2.06 |
|  | 4 | *EF1* | 1.3 | *EF1* | 1.14 | *EF1* | 1.25 | *EF1* | 2.66 | *EF1* | 4 |
|  | 5 | *His H3* | 3.33 | *GAPDH* | 2.28 | *His H3* | 2.16 | *His H3* | 4.05 | *His H3* | 5.23 |
|  | 6 | *GAPDH* | 5.28 | *His H3* | 2.39 | *GAPDH* | 3.29 | *GAPDH* | 5.55 | *GAPDH* | 5.73 |

Across most tissues analyzed as well as in the combined dataset *18S rRNA, RPL5, ACTB*, and *EF1* were identified as the most stable reference genes, depending on tissue type. In contrast, *GAPDH* and *His H3* consistently exhibited higher SV, indicating greater expression variability. A notable exception was observed in head tissue, where *GAPDH* (SV = 0.40) was ranked as the most stable reference gene. Minor differences in gene ranking between NormFinder and geNorm were observed, as expected due to differences in the underlying statistical models.

### 3.5. Expression Stability Analysis by BestKeeper

The BestKeeper analysis, based on the standard deviation (SD) and coefficient of variation (CV) of Cq values (Andersen et al., 2004), was used to evaluate the expression stability of candidate reference genes. Lower SD values indicate more stable expression; in accordance with the recommendations of Andersen *et al*. (2004), a SD value < 1 was considered an acceptability criterion for reference gene stability.

Across most tissues analyzed as well as in the combined dataset, genes identified as stable by geNorm and NormFinder also exhibited SD values < 1 according to BestKeeper. However, some discrepancies in gene ranking were observed. In particular, *GAPDH* showed acceptable expression stability in abdominal tissue according to BestKeeper (SD = 0.72), whereas it was ranked among the least stable genes by geNorm, NormFinder and ΔCt method for this body part Conversely, *EF1*, identified as stable in head tissue by geNorm, NormFinder and ΔCt method, displayed an SD value > 1 according to BestKeeper, indicating increased expression variability based on this algorithm (Table 2 and figure 4).

In contrast, *His H3* consistently exceeded the recommended variability threshold in BestKeeper, indicating unstable expression across all tissues analyzed as well as in the combined dataset. When all tissues were analyzed together, the ranking obtained with BestKeeper differed slightly from those obtained with NormFinder, geNorm and ΔCt method; nevertheless, *His H3* and *GAPDH* were concordantly identified as the least stable candidate reference genes (Table 2 and figure 4).

### 3.6. Expression stability analysis using the ΔCt method

The ΔCt method was used to evaluate the expression stability of candidate reference genes based on the mean and standard deviation (SD) of Cqvalues for each gene. Lower SD values indicate more stable expression. The ranking of reference genes obtained using the ΔCt method was largely consistent with that provided by the NormFinder algorithm, with only minor differences observed (Table 2, Figure 4).

Across most tissues analyzed as well as in the combined dataset, *RPL5* was identified as the most stable reference gene for the abdomen, legs, and the combined dataset, *GAPDH* for the head, and *EF1* for the whole body (Table 2). In contrast, *His H3* and *GAPDH* generally exhibited higher SD values, indicating increased expression variability. A notable exception was observed in head tissue, where *GAPDH* showed stable expression.

The ΔCt method further confirms the presence of body partsdependent variations in reference gene expression stability and highlights the need for tissue-specific validation prior to RT-qPCR data normalization, in agreement with the results obtained using the geNorm, NormFinder, and BestKeeper methods and in compliance with the MIQE 2.0 recommendations (Bustin et al., 2025).

### 3.7. Global analysis of candidate reference gene stability

The four statistical algorithms (geNorm, NormFinder, BestKeeper, and the ΔCt method) generated distinct ranking profiles. To obtain a consensus result, a comprehensive analysis based on the RefFinder tool was performed, enabling an integrated ranking of the candidate reference genes (Xie et al., 2012). RefFinder calculates the geometric mean of the rankings derived from the different methods; lower values indicate more stable gene expression (W. Wang et al., 2023). Considering the optimal number of reference genes, the most suitable genes were identified for four tissue types as well as for the combined dataset.

For abdominal tissue, *18S rRNA*, followed by *RPL5*, *ACTB*, and *EF1*, was ranked as the most stable reference gene (Table 2 and figure 4). In the head, *ACTB* showed the highest stability, followed by *GAPDH*, *18S rRNA*, and *RPL5*. For legs, the stability order was *RPL5, ACTB, 18S rRNA*, and *EF1*. For whole-body samples, *EF1* was identified as the most stable reference gene, followed by *18S rRNA, RPL5*, and *ACTB*. When all samples were analyzed together, *18S rRNA*, followed by *RPL5, ACTB*, and *EF1*, exhibited the highest overall stability (Table 2 and figure 4).

In contrast, *His H3* and *GAPDH* generally showed lower expression stability across most of the analyzed tissues as well as in the combined dataset. A notable exception was observed in head tissue, where *GAPDH* exhibited stable expression.

The Venn diagram (figure 5) shows that *ACTB, RPL5*, and *18S rRNA* are consistently identified as stable reference genes across all tested tissues, as well as in the integrated analysis of all tissues. This convergence indicates that these genes exhibit robust and consistent expression stability regardless of the tissue analyzed, making them suitable candidates for global normalization of RT-qPCR data. In contrast, *GAPDH* was identified as a tissue-specific reference gene, showing stable expression exclusively in head tissue.

**Figure 5:**
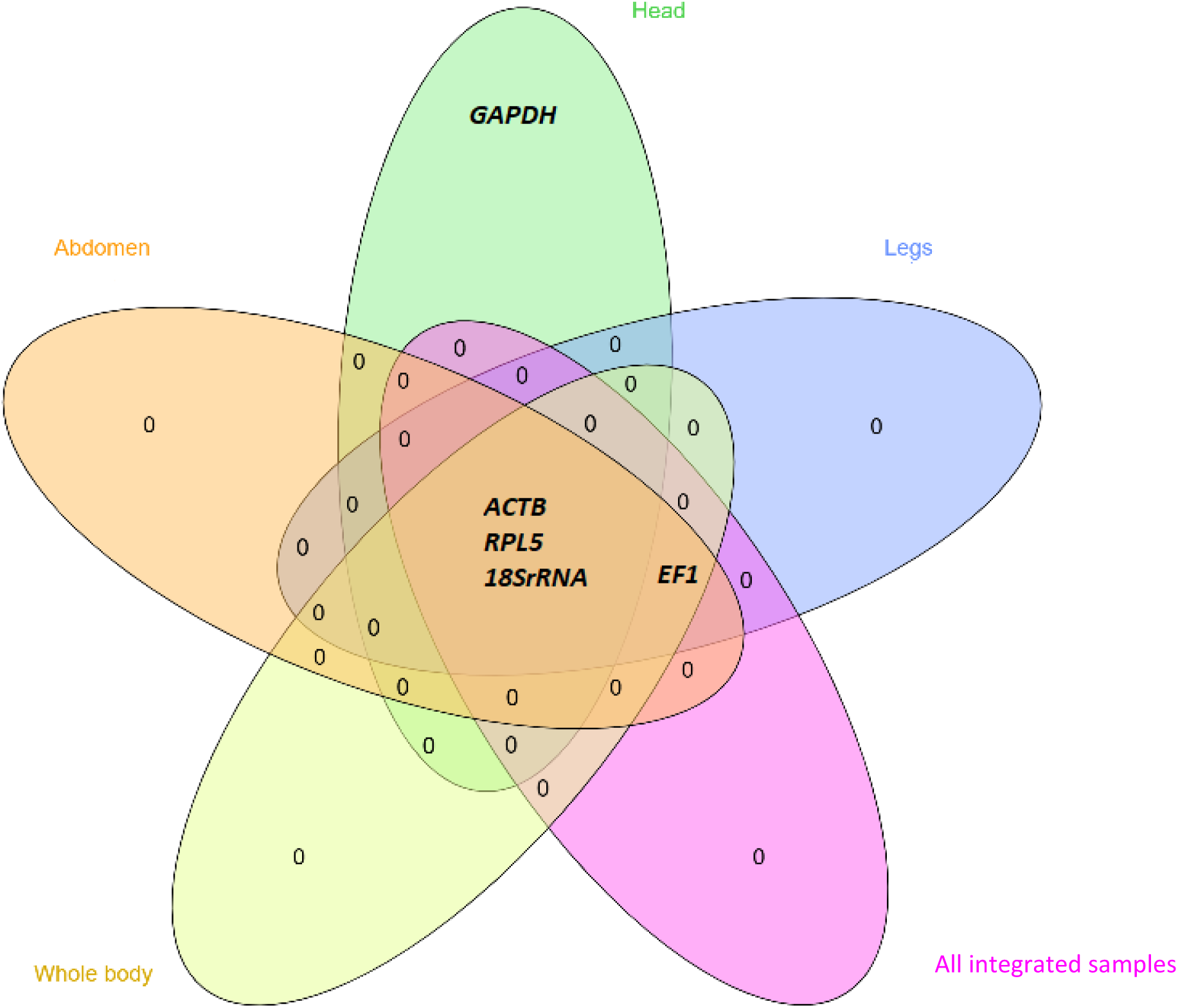
Venn diagram showing the number of genes and the overlap among the most stable genes in the different tissues of tropical cricket (*Gryllodes sigillatus*), as determined by the RefFinder analysis.

### 3.8. Pairwise Variation Analysis of Candidate Reference Genes Using R

The optimal number of reference genes required for accurate normalization of each tissue type and for obtaining reliable RT-qPCR results was determined using pairwise variation analysis (Vn/Vn+1) implemented in R. In accordance with the recommendations of Vandesompele and collaborators a Vn/Vn+1 value below 0.15 indicates that the inclusion of an additional reference gene does not significantly improve normalization quality (Hellemans & Vandesompele, 2014; Vandesompele et al., 2002).

For abdominal tissue, the V2/V3 value was 0.14. Although this value was slightly higher than the V3/V4 and V4/V5 values, it remained below the recommended threshold of 0.15. These results indicate that a combination of two reference genes is sufficient to ensure reliable normalization of target gene expression in the abdominal tissue of *G. sigillatus* (Figure 6) Similarly, pairwise variation analysis showed that V2/V3 values were below the 0.15 threshold for head, leg, and whole-body samples, suggesting that two reference genes are also sufficient for gene expression normalization under these experimental conditions (Figure 6).

**Figure 6:**
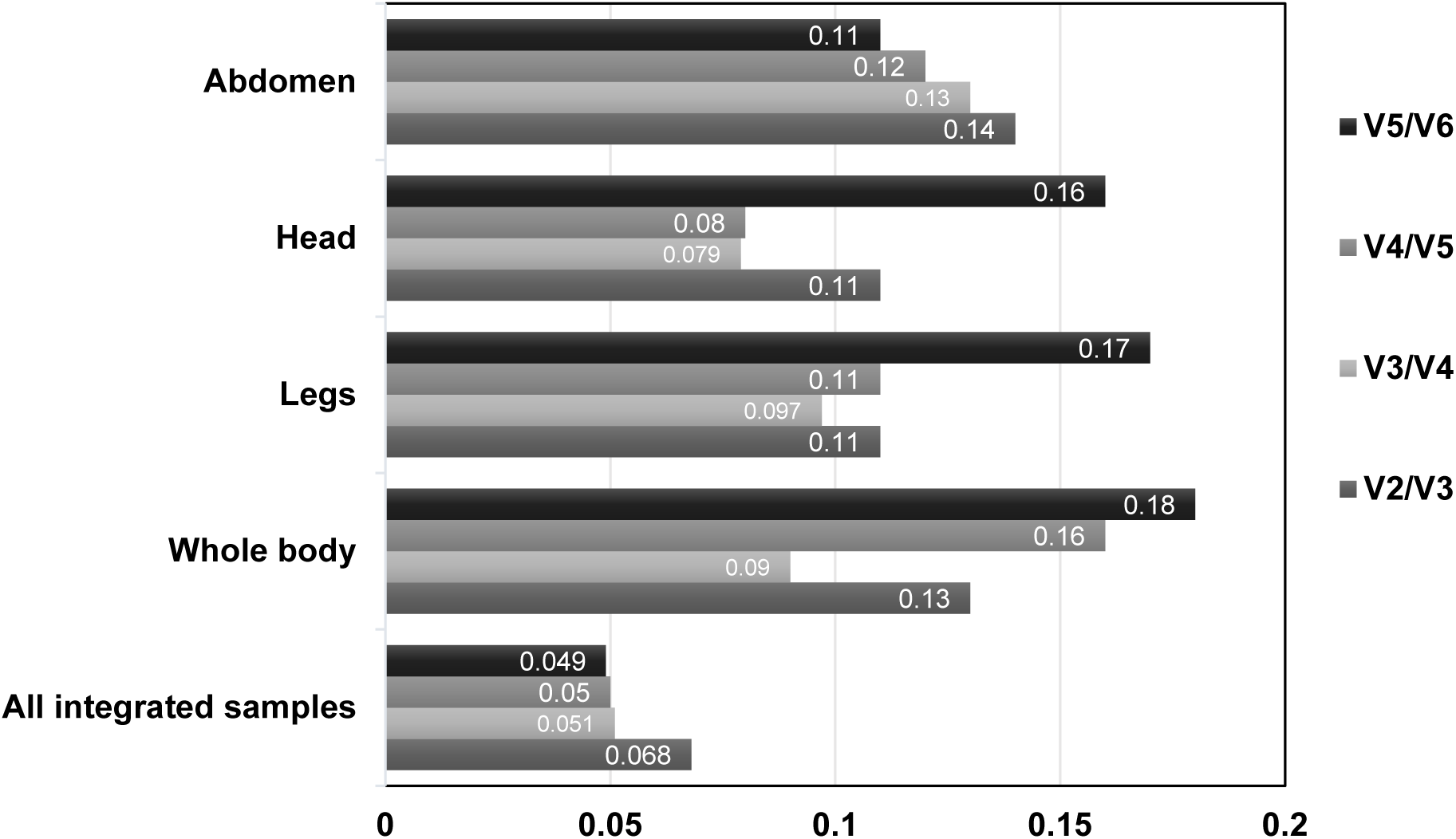
Pairwise variation (Vₙ/Vₙ₊₁) for determining the optimal number of reference genes. A value lower than 0.15 indicates that adding an additional reference gene (n+1) does not significantly improve the normalization factor and is therefore not required to ensure reliable gene expression normalization.

When all samples were combined, the V2/V3 value was higher than the V3/V4 and V4/V5 values; however, it remained well below the critical threshold, with a value of 0.068. This observation confirms that the use of two reference genes represents the optimal normalization factor for global gene expression analysis (Figure 6). Overall, these results demonstrate that the inclusion of a third reference gene does not provide a significant improvement in normalization, as evidenced by the absence of a substantial reduction in pairwise variation.

### 3.9. Verification of Reference Genes

To analyze the relative expression associated with candidate reference genes identified as the most stable and the most unstable in each tissue, we used the *EF2* gene, which is stably expressed in insects. The relative expression of *EF2* was used to verify the stability and applicability of the reference genes.

Based on the analysis of the results, we observed that in head tissue, the relative expression values of the target gene differed depending on the normalization strategy used. Normalization using unstable reference genes (*His H3* and *EF1*) resulted in significantly higher relative expression values compared with normalization using stable reference genes (*ACTB* and *GAPDH*) (paired two-tailed t-test, *p* = 0.047 < 0.05), with a large effect size (partial η² = 0.36) (Figure 7).

**Figure 7:**
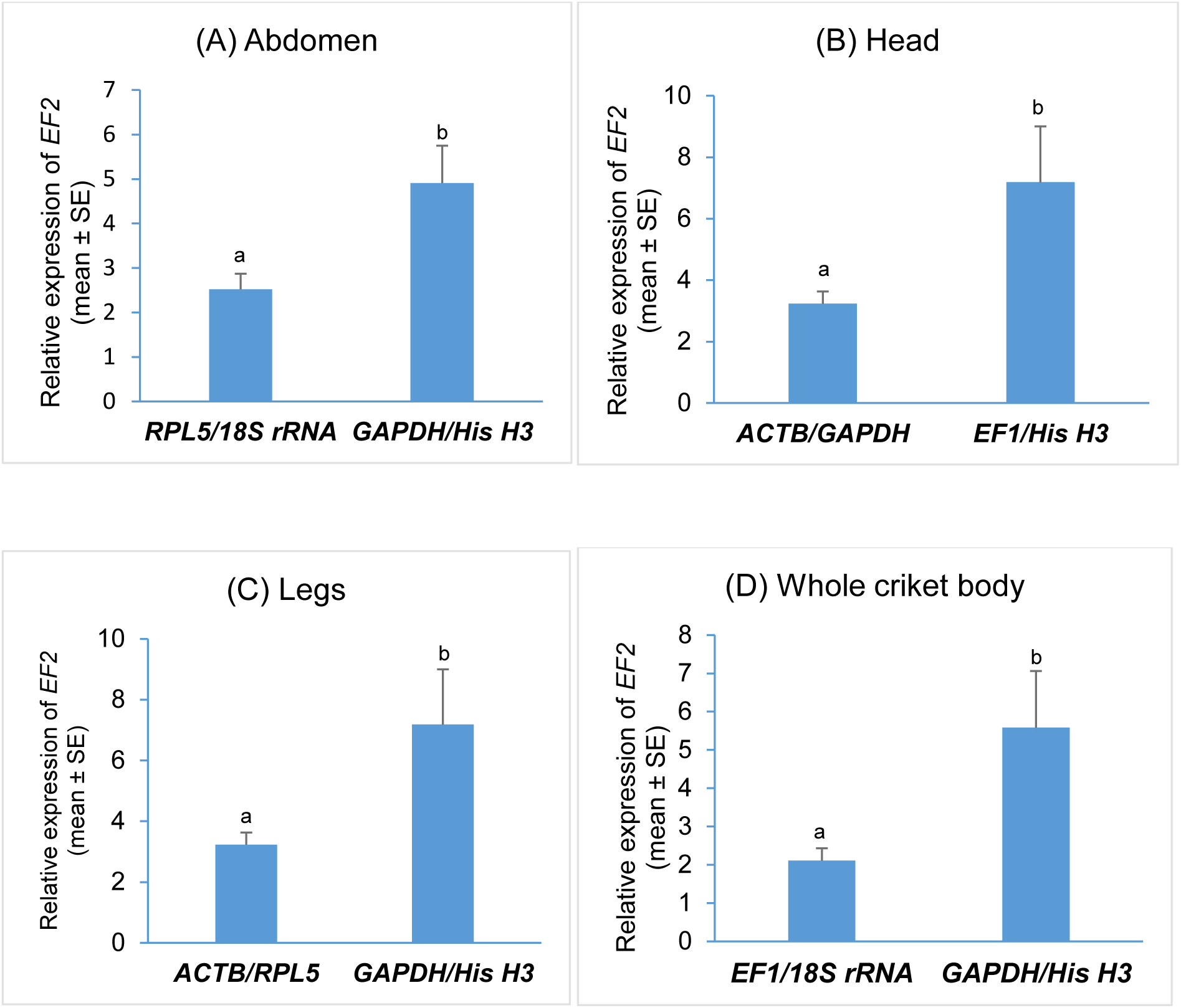
Comparison of *EF2* gene expression levels among four tissues of *Gryllodes sigillatus*. Relative expression was normalized using the two most stable reference genes and the two least stable (unstable) reference genes and calculated using the 2-ΔΔCq method. Statistical comparisons were performed using paired two-tailed Student’s t-tests, with P < 0.05 considered statistically significant. Data are presented as mean ± SE. Different letters indicate statistically significant differences among tissues (P<0.05).

Similar results were observed in the other tissues analyzed. In these tissues, the use of less stablereference genes (*His H3* and *GAPDH*) also led to significantly higher relative expression values compared with the corresponding stable reference genes, with statistically significant differences (*p* < 0.05) (Figure 7).

## 4. Discussion

Due to its high sensitivity, specificity, precision, and reproducibility, RT-qPCR has established itself as the gold standard method for quantitative analysis of gene expression, largely surpassing conventional approaches such as Northern hybridization or semi-quantitative PCR (Bustin et al., 2009; Wong & Medrano, 2005). It is now an essential tool for the detection and quantification of nucleic acids in a wide variety of experimental contexts ((Bustin et al., 2013; Hellemans & Vandesompele, 2014). However, the robustness and reliability of RT-qPCR results critically depend on the choice of reference genes used for normalization (Gutierrez et al., 2008; Hellemans et al., 2007; Vandesompele et al., 2002). Contrary to the long-accepted assumption of constitutive and stable expression, numerous studies have demonstrated that no reference gene is universally stable, and that the same gene may exhibit significant expression variations depending on the species, body parts, or experimental conditions (Han et al., 2021; W. Kong et al., 2024; Zhai et al., 2014). The inappropriate use of an unstable reference gene, or reliance on a single gene for normalization, can therefore introduce significant bias, mask true biological variations, or generate interpretative artifacts. These limitations highlight the need for rigorous and context-specific selection of reference genes to ensure reliable quantification of target gene expression (Bustin et al., 2009; Q.-Y. Li et al., 2019). Although several studies have validated reference genes in *Hemiptera*, *Lepidoptera*, *Coleoptera*, and *Diptera* (Lü et al., 2018), data remain insufficient for certain species, thereby justifying the need for dedicated validation analyses.

In insects, genes commonly used as reference genes are generally involved in fundamental cellular functions. Among them, beta actin (*ACTB*) encodes a major structural protein of the cytoskeleton and is expressed, at variable levels, in many cell types (Lü et al. 2018). Due to its central role, it is often considered an ideal reference gene for RT-qPCR analyses and is among the most frequently studied genes in the literature (Fu et al., 2013; Han et al., 2021; Zhang et al., 2015).

In addition, 18S ribosomal RNA and genes encoding ribosomal proteins (RP) were selected as candidates in the present study because of their essential structural role and their strong evolutionary conservation (Fu et al., 2013; Yuan et al., 2014). As the most abundant ribosomal component in eukaryotes, *18S rRNA* is frequently used as an internal reference gene in gene expression analyses (Lü et al., 2018). Similarly, genes of the *RPL* family, which represent approximately 18.55% of the reference genes reported in insect studies, have been among the most widely selected in recent years (Lü et al., 2018). Most studies highlight the relative stability of their expression across different tissues and experimental conditions (W. Kong et al., 2024; C. Yang et al., 2015).

Likewise, glyceraldehyde-3-phosphate dehydrogenase (*GAPDH*), a key enzyme of glycolysis, is widely used as a reference gene in insects (W. Zhao et al., 2022). It has been identified as a stable reference gene across developmental stages in *Rhopalosiphum padi* (M. Li et al., 2021), as well as a tissue-specific reference gene in *Lymantria dispar* (Yin et al., 2020), which motivated its inclusion among the candidate genes evaluated in the present study.

Elongation factor 1 alpha (*EF1*) is a protein involved in the delivery of aminoacyl-tRNAs to the ribosome during translation (Roger et al., 1999). Because of its central role in protein synthesis, *EF1* is frequently considered a reliable reference gene and is also among the most commonly used genes in insect phylogenetics (W. Kong et al., 2024; Liang et al., 2014).

Histone H3 genes in insects encode proteins essential for nucleosome structure and exhibit a high degree of evolutionary conservation (Delaney et al., 2018). This conservation, combined with their fundamental structural role in chromatin organization, justifies their frequent use as molecular markers in phylogenetic studies and also supports their selection as candidate reference genes for the normalization of gene expression data by RT-qPCR (Delaney et al., 2018; Elsaesser et al., 2010), provided that their expression stability is rigorously validated. These elements justify their inclusion as candidate genes in our stability analysis, in order to evaluate their suitability for RT-qPCR data normalization in *G. sigillatus*.

To limit biases associated with the use of a single analytical tool, four complementary approaches geNorm, NormFinder, BestKeeper, and the ΔCt method were applied to establish stability rankings of the six candidate reference genes in *G. sigillatus* across four distinct tissues. The rankings obtained revealed slight variations among the different analytical approaches (Table 2).

In the abdomen, NormFinder and the ΔCt method identified *RPL5, 18S rRNA, EF1*, and *ACTB* as the genes showing the greatest expression stability. In contrast, geNorm proposed a slightly different hierarchy, ranking *18S rRNA, ACTB, EF1*, and *RPL5* as the optimal combination of stable genes. BestKeeper, however, also ranked *GAPDH* among the stable genes in this tissue, whereas this gene was identified as one of the least stable by the other algorithms, along with *His H3*, which was consistently classified among the least stable genes by all algorithms. This type of divergence, particularly between BestKeeper and the other algorithms, has already been reported in studies by Kolsher et al. (2020) and Kaloužná et al. (2017) (Kałużna et al., 2017; Köhsler et al., 2020). These differences may be attributed to the methodological specificities of BestKeeper, which appears to be less suitable for comparing reference genes exhibiting highly contrasting expression levels. Indeed, when expression values differ markedly, the parametric correlations used by this algorithm may lead to divergent rankings, which could explain the position assigned to *GAPDH* (Pfaffl et al., 2004).

In the head tissue, BestKeeper and geNorm provided an identical ranking of the most stable candidate genes, namely *ACTB, 18S rRNA, GAPDH*, and *RPL5*. Although NormFinder and the ΔCt method showed slight variation in the order of the three top-ranked genes, the ranking of the other candidates remained similar. Comparable trends were observed for the other tissues, as well as for the integrated analysis of all tissues, where the rankings were either consistent or characterized by minor divergences. These moderate differences are consistent with results reported in the literature and can be explained by the fact that these tools are based on distinct algorithmic principles, which may lead to slightly different rankings of expression stability (Shi et al., 2013; J. Tang et al., 2022; Xue et al., 2024).

In this context, an integrated analysis was performed using the online tool RefFinder, which combines the results of the four algorithms. This combined approach allowed the establishment of a final ranking of the reference genes. The results reveal that the stability of the candidate reference genes varies according to the tissues analyzed as well as in the overall analysis integrating all tissues. Nevertheless, *18S rRNA, RPL5, ACTB*, and *EF1* were identified as the most stable genes in the abdomen, legs, whole body, and in the integrated analysis of all tissues. In contrast, in head tissue, a distinct stability profile was observed, with *ACTB, GAPDH, 18S rRNA,* and *RPL5* identified as the most stable reference genes. This variation in ranking according to tissue type is confirmed by the study of Sellamuthu et al. (2022), which highlighted different classifications of candidate reference gene stability depending on the body parts analyzed in the Eurasian spruce bark beetle *Ips typographus* (Sellamuthu et al., 2022). In addition, the study by Yang et al. (2025) shows that *ACTB* is stable in the legs and wings but unstable in the abdomen and thorax, while, in the same study, *EF1* is stable in head and antennal tissues and unstable in the abdomen and thorax in *Scotogramma trifolii*, which confirms our differences in ranking (A. Yang et al., 2025).

Recent studies have also identified gene encoding ribosomal proteins such as *RPL5* as stable reference genes for RT-qPCR in *Diuraphis noxia* (Sinha & Smith, 2014), which is consistent with our results. Our results show that *GAPDH* is unstable in all tissues except the head, which is similar to the study conducted in *Meliponini* (Freitas et al., 2019), where this gene shows instability across different tissues. In contrast, in the study by Yang et al. (2025) on *Scotogramma trifolii*, *GAPDH* is described as stable, but with variable ranking depending on tissue type (A. Yang et al., 2025).

Furthermore, several studies conducted in *Apis mellifera* (Jeon et al., 2020), the whitefly *Dialeurodes citri* (W. Kong et al., 2024), and *Athetis dissimilis* (J. Tang et al., 2022) show that *18S rRNA* is stable in the different tissues studied, with variations in ranking depending on tissue type.

Finally, regarding the gene identified as the most unstable across all tissues analyzed, *HisH3*, this instability has also been reported in *Locusta migratoria* across the different tissues examined (Q. Yang et al., 2014).

The number of reference genes required for normalization was determined using analyses performed with the R software, based on the calculation of pairwise variation (Vn/n+1) in accordance with the approach proposed by Vandesompele et al. (2002) (Vandesompele et al., 2002). With the aim of optimizing the robustness and precision of RT-qPCR data, the use of multiple internal reference genes is recommended, with the choice of the appropriate number generally relying on a variation threshold set at 0.15 (Hellemans & Vandesompele, 2014; Shu et al., 2018; Vandesompele et al., 2002). In the present study, V2/V3 values were below this threshold, indicating that the use of two reference genes allows adequate normalization. These observations are consistent with previous studies showing that the use of a single reference gene in RT-qPCR analyses can generate notable bias in the normalization process (Guo et al., 2020; W. Kong et al., 2024; C. Yang et al., 2015).

In conclusion, these results highlight the central role of rigorous reference gene selection in obtaining reliable and interpretable RT-qPCR data (Bustin et al., 2013, 2025; Hellemans & Vandesompele, 2014). Conversely, normalization based on inappropriate or variably expressed reference genes is likely to bias the analysis and compromise the correct interpretation of target gene expression (Guo et al., 2020; Morammazi & Shokrollahi, 2020).

However, this study has certain limitations, including the evaluation of a restricted number of candidate reference genes and experimental conditions. Nevertheless, it represents an initial step toward the identification and validation of reference genes in these species. Future studies should expand this validation to different developmental stages, as well as to stress and infection conditions, in order to strengthen the robustness of reference gene selection. Moreover, integrating these validated reference genes into RT-qPCR-based diagnostic assays could support the development of reliable molecular tools for monitoring insect health and improving biosafety in insect production systems.

## 5. Author’s contributions

HBM: Conceptualization, Experimentation, Formal analysis, Writing – original draft, Writing – review & editing. FR: Experimentation, Formal analysis, Writing – review & editing. RB: Conceptualization and Experimentation. FM and MOBB: Project administration, Conceptualization, Validation, Formal analysis, Writing – review & editing, Funding acquisition, Supervision.

